# Estimating plant abundance in irregular polygons: plot-based sampling with a conditional spatial configuration

**DOI:** 10.1101/2020.08.17.253864

**Authors:** Christian Damgaard

## Abstract

When estimating plant species abundance using plot-based methods in irregular polygons, there are some constraints to the spatial configuration of the sampling plots: i) the sample plots must be positioned within the polygon, ii) the sample plots may not overlap, iii) due to the spatial variation in abundance often found in natural plant communities, the spatial coverage must be relatively large in order to decrease the sampling variance, iv) due to a possible mosaic structure of the vegetation, systematic designs should be avoided due to the possibility of biased estimates. Here, we suggest a protocol using simulations of the possible random spatial configurations to select a conditional spatial configuration that meets these constraints.

## Introduction

It is a common objective in plant ecology to compare the vegetation among different sites or treatments, and an important characteristic of the vegetation that is suitable for comparison is the abundances of the different plant species. A number of different measures are used for quantifying plant abundance, i.e. probability of occurrence, cover, density, vertical density and biomass (Damgaard 2014), and these abundance measures rely on a plot-based sampling scheme, where only the abundance of the vegetation within relatively small plots are measured (Kenkel et al. 1989, Kent and Coker 1992).

Natural plant communities may be dominated by plant species where it is difficult to distinguish individual plants, e.g. grasses, and most species show some form of spatial aggregation. Such key features of plant communities in combination with the type of research question play an important role when deciding which plant abundance measure to apply (Kenkel et al. 1989). For example, if a possible target species is relatively rare and evenly distributed, then the probability of occurrence in relatively large plots may be preferred, whereas if the target species is dominant and spatially aggregated, then cover in relatively small plots may be a more suitable measure of abundance. Furthermore, the different abundance measures have different statistical sampling properties, and if plant species are spatially aggregated, then more plots are needed to obtain the same precision as spatially dispersed plant species (Damgaard 2013).

When the method of measuring abundance, plot size and sampling intensity has been decided, then the spatial configuration of the sampling plots must be considered. Generally, when the probability of positioning a sampling plot at a specific place is independent of the vegetation in the plot, then the sampling mean of species abundance is an unbiased estimate of the population mean (Horvitz and Thompson 1952). Furthermore, if there is spatial variance in the vegetation, sampling designs that ensure the sample is well-distributed over the study region will, in general, improve estimation of population abundance relative to designs that do not (Munholland and Borkowski 1996, Borkowski 2003). Since almost all natural plant communities have some degree of spatial variation, it has therefore been a common practice among empirical plant ecologists to increase the spatial coverage of the sample plots by using a systematic sampling design, where the plots are arranged in a regular grid or equidistant along transects (e.g. Cochran 1977). However, systematic designs may lead to biased estimates in a mosaic vegetation if the distance between sampling plots is similar to the spatial scale of the mosaic structure of the vegetation. This bias arises because when the spatial scales of the sampling plots and the mosaic are similar, then the probability of sampling a plot is no longer independent of the vegetation (Fig. 1, Munholland and Borkowski 1996, Borkowski 2003). Since the spatial scale of different mosaic structures of the vegetation typically is unknown in the design-planning phase and varies among plant species, a systematic design of sampling plots is problematic. Consequently, it has been suggested to replace both random and systematic designs with Latin square sampling approaches, which provide unbiased estimates and where the sampling coverage is increased relative to a random design (Munholland and Borkowski 1996, Borkowski 2003, Salehi 2006). However, Latin square sampling is not suited for the sampling of relatively small plots in irregular polygons.

**Fig. 1.**
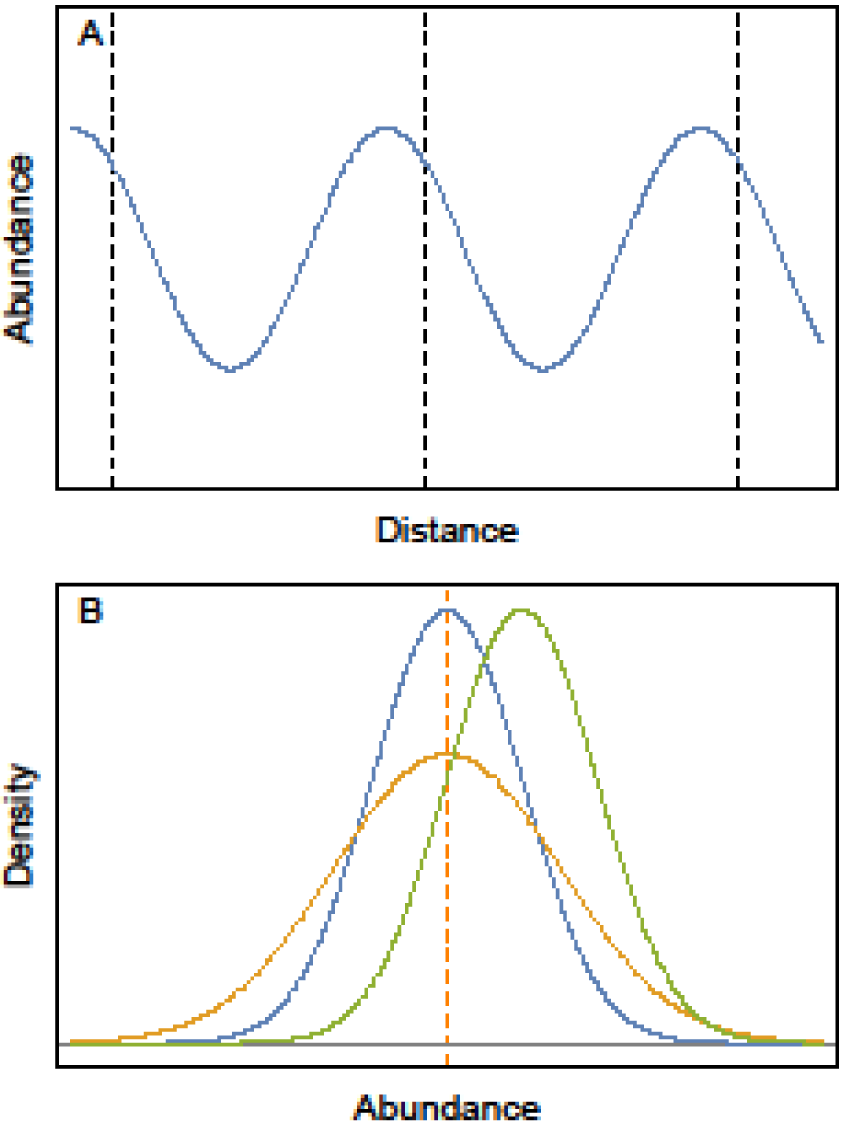
Conceptual figures of the effect of sampling design on the estimated abundance. A. If the scale of a systematic sampling design (dashed lines) is similar to the scale of the vegetation mosaic (the full line shows the oscillating abundance of a single species), then the estimate of abundance may be biased. B. Expected distribution of sampling mean at three different sampling designs: simple random sampling (orange line), random sampling with increased spatial coverage (blue line) and systematic sampling (green line). The population mean is the dashed orange line.

A trivial, but important feature of plot-based sampling is that the size of the plots must be taken into account when the spatial configuration of the sampling plots is laid out. More specifically, the entire plot must be placed within the borders of the polygon, and the plots should not overlap. Thus, two requirements for the spatial configuration of the sampling plots must be fulfilled: i) the center of each plot must be at least half the plot size distance (the radius in a circle, or half the length of the side in a square) from the polygon border, and ii) in order that the plots do not overlap, the distance between the center of two plots must be at least equal to the plot size distance.

Here, we suggest selecting the spatial configuration of sample plots as the conditional spatial configuration that best fulfills the above-mentioned requirements out of many simulated random spatial configurations.

## Selecting a conditional spatial configuration

The joint distribution of the distances between *n* points in the plane may be found explicitly (Molchanov 2012). However, since there is no closed solution for points within an irregular polygon, we used simulations to determine the geographic positions of sampling plots in an irregular polygon. In a simulation, a fixed number of sampling plots are positioned randomly in the polygon to provide a realization of the spatial configuration. This simulation is repeated many times, and for each simulated spatial configuration, the minimum distance from the plots to the border of the polygon and between all plots is determined.

From all the simulated sets of spatial configurations, we selected the intersection of the sets that fulfilled both selection criteria, i.e. the minimum distances should be sufficiently large, so that the entire plot was placed within the borders of the polygon and the plots did not overlap.

The selection criteria may be set either as the upper 97.5% percentile of the simulated minimum distances, or as a fixed minimum distance. If the area of the polygon is relatively large compared to the number of plots, then it is advantageous to set the criterion for the minimum distance to the polygon border as half the plot size distance and restrict the criterion for the minimum distance among the plots using e.g. the upper 99% percentiles of the simulated minimum distances.

The final conditional spatial configuration of the sample plots was determined by selecting the spatial configuration with the best spatial coverage of the sampling plots. This selection was performed by a visual inspection of the remaining spatial configurations in the intersection set.

Note that the above-mentioned selection processes are done at the computer using GIS software with only a general knowledge of the vegetation type and treatment configuration. Consequently, the different plant species and their associations have the same probability of being included in one of the selected sample plots, and the selected conditional spatial configuration will, consequently, provide an unbiased estimate of species abundance (Horvitz and Thompson 1952).

A Mathematica notebook (Wolfram 2020) that uses a polygon constructed with QGIS was used to generate the limited set of possible spatial configurations. The notebook is included as supplementary material.

## Results

The above-mentioned selection procedure is illustrated using an example of an irregular polygon (Fig. 2), where 10,000 realizations of 50 random sampling plots were simulated. Histograms of the simulated minimum distances from the plots to the border of the polygon and between all plots are shown in Fig. 3. The mean minimum distance from the border of the polygon was 1.43 m and the 97.5% percentile of the minimum distances was 5.16 m, whereas the mean minimum distance between plots was 5.28 m and the 97.5% percentile of the minimum distances was 11.63 m.

**Fig. 2.**
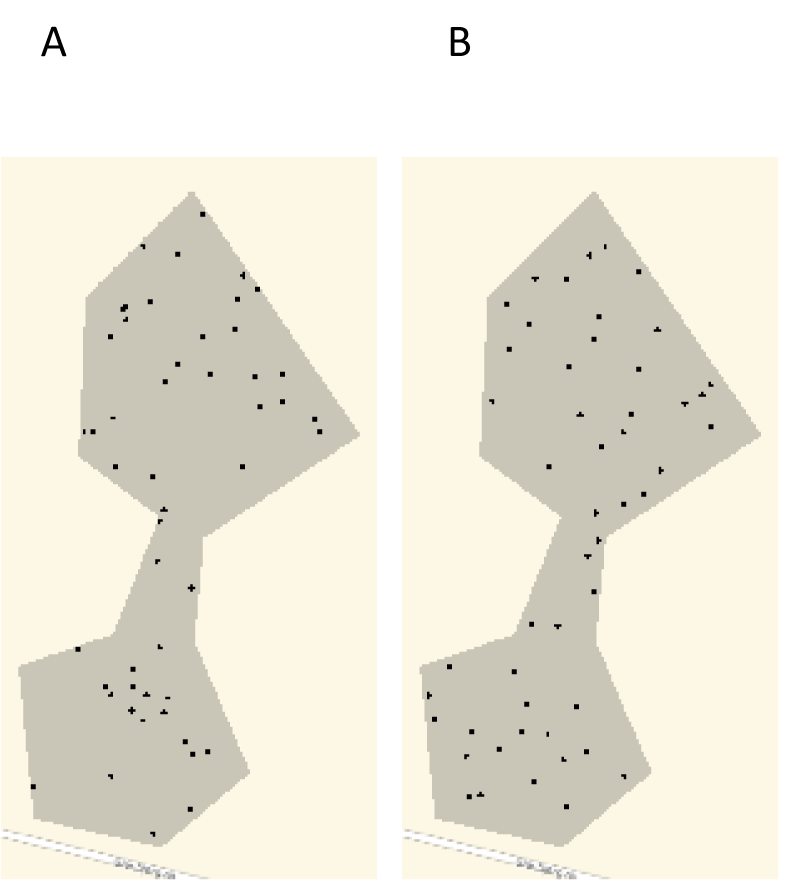
An irregular polygon of 132,781 m2 with 50 randomly positioned sampling plots (A) or 50 sampling plots in a conditional spatial configuration (B).

**Fig. 3.**
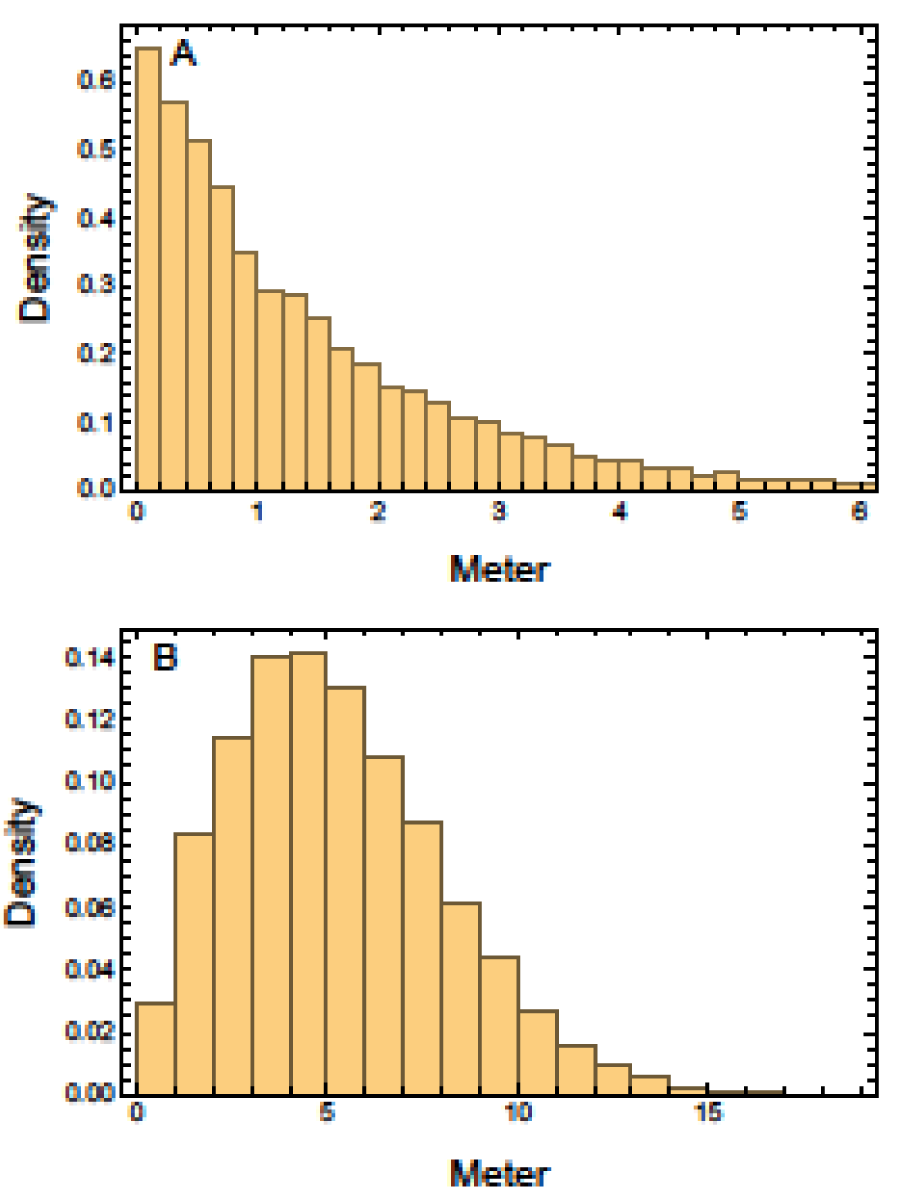
The smallest distance between the border and a plot (A) and between two plots (B) in 10,000 samples of 50 random sampling plots in the irregular polygon shown in Fig. 2.

In this particular case, the 97.5% percentile selection criteria of the minimum distances was appropriate for measuring the probability of occurrence in circle plots with a radius of 5 m, as performed in the Danish monitoring program of terrestrial habitats (Nielsen et al. 2012).

After using the 97.5% percentile selection criteria, the interaction set had five spatial configurations and the final conditional spatial configuration was selected as the one with the best spatial coverage after a visual inspection.

A simulated realization of the spatial configuration of 50 random points may be compared with the selected conditional spatial configuration (Fig. 2), where distance requirements are fulfilled for circle plots with a radius of 5 m and the spatial coverage has been increased considerably.

## Discussion

When estimating plant species abundance using plot-based methods in irregular polygons, there are some constraints to the spatial configuration of the sampling plots: i) the sample plots have must be positioned within the polygon, ii) the sample plots may not overlap, iii) due to the spatial variation in abundance often found in natural plant communities, the spatial coverage must be relatively large in order to decrease the sampling variance (Munholland and Borkowski 1996), iv) due to a possible mosaic structure of the vegetation, systematic designs should be avoided due to the possibility of biased estimates (Munholland and Borkowski 1996). All these requirements are met with the suggested method of selecting a conditional spatial configuration of sample plots. However, if the area polygon is relatively small compared to the number of plots, then it may not be possible to meet the selection criteria and then the plot size or the number of sampling plots must be reduced.

Unlike when using a systematic design, the conditional spatial configuration of sample plots ensures variation among plot distances, which enables us to estimate the spatial scale or grain size of the possible mosaic structure of the vegetation using e.g. semi-variograms.

In most cases, it is recommended to examine the effect of the spatial variation of the vegetation on the relevant statistical inference, e.g. using the stochastic partial differential equations approach for modeling continuous spatial processes with a Matérn covariance, which has been implemented using the integrated nested Laplace approximation in the R-INLA package (Blangiardo and Cameletti 2015, Krainski et al. 2018).

The final selection step, where the spatial configuration with the best spatial coverage was selected, was performed by a visual inspection. This procedure is unproblematic in practice, since it is always possible to set sufficiently limiting selection criteria so that only a few spatial configurations remain in the intersection set. However, it may be advantageous in some cases to develop an automated approach of maximizing the spatial coverage using objective measures of the degree of spatial coverage.

Munholland and Borkowski (1996) discuss the estimation of the sampling variance due to the spatial configuration of the sample in vegetation plots with spatial variation and aggregation using a space filling approach, but their suggestion is not suitable when only relatively small plots are sampled from a larger area.

To our knowledge, the importance of the sampling bias when applying a systematic design (Munholland and Borkowski 1996, Borkowski 2003) has not been investigated in natural plant communities. However, since it is relatively easy with modern handheld GPS to avoid the possible bias, it seems most prudent to avoid systematic sampling design when studying natural plant communities with mosaic vegetation patterns.

In some plant ecological studies, gradient directed transects (Gillison and Brewer 1985), or so-called “stratified sampling methods”, where the area of different strata is unknown, are used to characterize the variation in the vegetation. However, these sampling methods violate the assumption that different plant species have the same probability of being included in one of the selected sample plots (Horvitz and Thompson 1952) or have an alternative statistical model, thus, these sampling methods cannot be used for estimating plant abundance.

Finally, it is worth mentioning that it is possible to estimate plant abundance using plot-less distance-based sampling procedures (e.g. Cochran 1977) and, more specifically, the probability of occurrence cover, and a combined feature of two abundance measures, integral occurrence probability may be measured using a plot-less distance-based sampling procedure (Van Calster and Damgaard 2017).

## Supporting information

Mathematica notebook

## Acknowledgements

I am grateful to Rikke Reisner Hansen, Morten Tune Strandberg and Peter Borgen Sørensen for input and valuable discussions.

